# Indole acetic acid and lipopeptide-producing endophytic bacteria from *Taxus chinesis*: toxicity evaluation of metabolic products

**DOI:** 10.1101/2020.06.08.139915

**Authors:** Mengmeng Dong, Bo Yuan, Jingyi Wang, Lizhong Fu, Guoyin Kai, Jihong Jiang

## Abstract

Endophytes play an important role in plant growth and development. Some one can produce auxins, ACCs, iron carriers, and so on to help plants grow and resist unhealthy growth environments. In addition, they can produce certain antimicrobial substances to resist pests and diseases. Among them, *Bacillus* is the most common beneficial endophytic bacterium in plants. In this paper, 20 IAA-producing strains were screened from endophytic bacteria isolated from *Taxus chinensis var. mairei* plant tissues by high-performance liquid-chromatography (HPLC). Based on the 20 IAA-producing strains, LC-TOF-MS technology was used to screen lipopeptide-producing *Bacillus* sp. As a result, three strains (KLBMPTC01, KLBMPTC10, and KLBMPTC29) of *Bacillus*-producing lipopeptides with abundant contents and species were obtained. According to the situation of the IAA and lipopeptides produced by these strains, KLBMPTC10 was selected as the experimental strain for later toxicological tests. In an Ames test and oral toxicity experiments in mice, we did not detect mutagenicity and other physiological toxicity. This is hoping to provide a theoretical basis for forest resource protection and biofertiliser production therewith.

## Introduction

An endophyte is an endosymbiont, often a bacterium or fungus(1), that lives within a plant for at least part of its life without causing apparent disease(2). In the long-term evolution, some endophytes co-evolve with the host and have a certain role in promoting the host. Some endophytes can promote the accumulation of plant medicinal active ingredients (3) and produce important active substances similar to host plants(4);And some endophytes can produce some substances such as indole-3-acetic acid (IAA) Siderophores, ACC deaminase to promote plant growth and development(5); even some endophytes such as the *Bacillus* can protect host from pests and diseases(6).

*Bacillus* sp. is the most common endophytic bacterium isolated from plant tissues and is a type of microbial population that is widely distributed in soil and plant (7). It plays an important role in plant growth, biological control, aquaculture, environmental protection, and medicine. Studies have shown that certain *Bacillus* can produce IAA to help plants grow(8) and promote the growth and disease-resistance of some aquatic economically-valuable animals (9, 10); Some *Bacillus* such as *Bacillus thuringiensis* and *Bacillus subtilis* were widely used as a microbial insecticide because they can produce some toxins to protect cash crops from pathogenic organism(11–13); Additionally, some *Bacillus* can be used to treat wastewater(14) and promote plant uptake of heavy metals in soil(15). Meanwhile, in medicine, some *Bacillus* sp. were acted as effective microorganisms in disease control and health improvement.

It is more common that *Bacillus* can produce IAA to promote plant growth(16, 17). IAA is one of the natural auxin in plants and it has an important position in plant growth and development(18), it can affect the elongation and splitting of plant cells(19). However, plant growth and development are inevitably accompanied by interference from disease. Coincidentally, some *Bacillus* sp. can produce some lipopeptide antimicrobial substances, such as surfactin, iturin, and fengycin, which can protect plants from the invasion by pathogenic bacteria (20).

Surfactin, iturin, and fengycin are the main lipopeptide active substances produced by *Bacillus* sp. Surfactants can dissolve and destroy cell membranes which display antiviral, antimycoplasma, and antibacterial activities(20). Iturin can quickly cause cell membrane damage, change their membrane permeabilisation properties, and allow leakage of intracellular substances which could explain why fungal spore germination and mycelial growth were inhibited. Fengycin exerts a significant disruptive effect on cell membranes, which can damage the phospholipid bilayer and lead to cell death(21–25). These three antimicrobial peptides have been widely used in agricultural production because of their strong resistance to plant pathogens.

Recently, most *Bacillus* with antimicrobial and growth-promoting properties are made into environmentally-friendly microbial agents and used instead of chemicals for fighting plant pathogens. However, while some Bacillus produce beneficial antibacterial substances, they can also produce substances harmful to human or animal health(26, 27). So, at the same time, human safety must not be overlooked. In this context, the present study focusses on the isolation of putative endophytic bacteria associated with *Taxus chinensis var. mairei* and evaluation of the traits of *Bacillus* isolates that produce IAA and lipopeptide antimicrobial substances. Furthermore, the toxicity of the endophytic *Bacillus* strain who can produce high IAA and peptide yields was studied in mice. We hope to provide a theoretical basis for forest resource protection and biofertiliser and bioenhancer production.

## Materials and methods

### Sampling and isolation of plant-associated bacteria

Healthy, natural *Taxus chinensis var. mairei* samples of root, stem and leaf were collected from the Lichuan in Hubei. The samples were surface-sterilised using previously described methods (28, 29), then washed for 2 min in 75% ethanol, then washed in sterile water, then washed for 15 min in 0.1% HgCl_2_, then washed in sterile water. The sterilised tissue samples were placed on NA agar and Luria Broth agar plates and incubated at 30 °C for 2 weeks to ensure the efficacy of surface sterilisation. The surface-sterilised samples were aseptically chopped into smaller fragments using a commercial blender and crushed using a sterile mortar and pestle, with sterile distilled water. Subsequently, 100 μL of the tissue extracts and serially diluted samples (10^-1^ to 10^-3^) were plated onto two different types of media. All plates were incubated at 30 °C for 2 weeks. Colonies were selected based on morphology. Selected colonies were transferred to fresh LB (5 g yeast, 10g NaCl, 10 g tryptone, 15 g agar, pH 7.0) to establish pure cultures of the bacteria.

### DNA extraction, sequencing, and analysis

The 16S rRNA gene sequences of the isolated strains were analysed to identify genus. The identities of the organisms were determined based on partial or nearly-full length 16S rRNA gene sequence analysis. The genomic DNA of each isolate was extracted using the method of Li et al. (30). The 16S rRNA genes from pure cultures were amplified using the primer pair 27F,1492R(27F:5′-CAGAGTTTGATCCTGGCT-3′,1492R:5′-AGGAGGTGATCCAGCCGCA-3′). PCR was conducted under the following conditions: initial denaturation at 95 °C for 5 min; 35 cycles at 95 °C for 30 s, 55 °C for 30 s, 72 °C for 90 s, followed by final extension at 72 °C for 10 min. The reaction mixture (50 μL) contained 25 μL of Taq PCR Master Mix, 2 μL of each primer at 5 μM, 19 μL of H_2_O, and 2 μL of genomic DNA. Subsequently, 5-μL PCR products were analysed electrophoretically in 1% (w/v) agarose gels stained with ethidium bromide. After visual inspection under UV light, 45-μL aliquots of PCR products were purified and sequenced at Shanghai Sangon Biological Engineering Technology & Services Co. Ltd. Sequences were then compared with16S rRNA gene sequences from reliable isolates listed in the EzBioCloud database (https://www.ezbiocloud.net/). A phylogenetic tree was constructed by the neighbour-joining method using the MEGA 7.0 software package.

### Fermentation, extraction, and evaluation of IAA-production ability

Each isolate was cultured in sucroseminimal liquid medium (glucose 1%, (NH_4_)_2_SO_4_ 0.1 %, K_2_HPO_4_ 0.2 %, MgSO_4_ 0.05 %, yeast extract 0.05 %, CaCO_3_ 0.05 %, NaCl 0.01 %, pH 7.2) supplemented with a final concentration 0.5 mg mL^-1^ of tryptophan at 30 °C and was shaken at 180 rpm in the dark). The 400 mL of liquid medium was inoculated with 4 mL of culture that had been grown overnight. After 5 to 10 days of cultivation, the fermentation pH of the extract was adjusted to 3.0 with concentrated hydrochloric acid(31, 32). Extraction with ethyl acetate was carried out as described by Singh et al. (33). The ethyl acetate extract was evaporated to dryness and the residue was dissolved in 2 mL of methanol. The samples were detected by using high-pressure liquid chromatography-time-of-flight mass spectrometry (HPLC-TOFMS) (34). All standard and sample solutions were passed through a 0.45-μm filter before use. The detection conditions were as follows: isocratic elution (methanol /water, 75:25, v/v) with Halo C18 column chromatography (4.6 × 150 mm, packed with 3.0 μm C18). The flow rate of the pump was 0.4 mL/min. The TOFMS conditions were as follows: under the electrospray ionisation ion source positive ion mode (ESI^+^), the ion source temperature was 550 °C, the voltage was 4500 V, the collision voltage was 70 eV, and the cluster voltage was 30 ev. The ratio *m/z* was set to 130 matching the target quantitative ion of indoleacetic acid. The detection limit was 0.0015-0.5388 μg/L, and the linearity was good over the range from 5-200 μg/L.

### Evaluation of lipopeptide-production ability of IAA-producting endophytic bacteria

The samples were dealt with as described in experiment “Fermentation, extraction, and evaluation of IAA-production ability”: separation and identification of lipopeptides in extracts were assayed by HPLC-TOFMS. The differences between lipopeptide compounds in strains were screened through comparison with a published database. The chromatographic conditions were as follows: gradient elution (acetonitrile/water, with Halo C18 column chromatography (4.6 × 150 mm, packed with 3.0 μm C18) and the gradient elution procedure was as follows: within 0-20 minutes, the proportion of acetonitrile decreased from 60% to 5.0%, and was kept for 10 minutes. Additionally, the flow rate of the pump was 0.5 mL/min. The TOF-MS conditions were as follows: under the electrospray ionisation ion source positive ion mode (ESI^+^), the ion source temperature was 400 °C, the collision voltage was 50 eV, and the cluster voltage was 10 ev; the scanning range was from100 ≤ *m/z* ≤ 2000.

### Bioassays: *in vivo* toxicity of KLBMPTC10 extraction in mice Preparation of KLBMPTC10 extract and in vivo toxicity in mice

The experiments were performed on ICR mice (female: male, 50:50) (weighing 18–22 g) obtained from the Xuzhou Medical College (Jiangsu, China). The animals were housed at 25 ± 5 °C at 60 ± 5 % relative humidity, and subjected to a 12 h light–dark cycle and left to acclimatise for 1 week before the experiments. They were fed standard laboratory chow and provided with water *ad libitum*. The experimental protocol was approved by the Local Animal Care Committee, and all the experimental procedures were carried out in accordance with the international guidelines for the care and use of laboratory animals (35, 36).

Strain KLBMPTC10 extraction was dealt with as described in experiment “Fermentation, extraction, and evaluation of IAA-production ability” and freeze-dried. Bacitracin was used as standard substance to determine the content of bacitracin in each sample by external standard method. Then the dried extract was dissolved in sterile distilled water and specimens with bacitracin at concentrations of 5, 10, 20, 50, and 100 μg/ml prepared for acute oral toxicity and Ames studies (37).

### Ames test

The test strains of TA97, TA98, TA100, and TA102 used in this study were all mutant strain of *Salmonella* which they cannot synthesize the necessary amino acid-histidine for standard bacterial culture (38). Before the experiment, the characteristics of the tested strains were identified to ensure that they met the requisite experimental standards. An Ames test was carried out according to the standard protocol without the S9 mix as described by Grúz et al. (39). Furthermore, 2-acetamidofluorene, Dexon, and sodium azide are three known positive mutagens with genotoxicity(40). And in this test, we used 20 μg/mL 2-acetamidofluorene, 50 μg/mL Dexon, and 20 μg/mL sodium azide as the positive control, the solvent group was DMSO and the experimental group is KLBMPTC10 bacterial extracts with concentrations of 5, 10, 20, and 50 μg/mL respectively. The blank control group was also set to observe spontaneous revertant counts. There were three replicates established of each treatment. Colonies were counted after 2 days of incubation at 37 °C using a bacterial colony counter (35).

### Acute oral toxicity study

To test the toxicity of strain KLBMPTC10, 36 mice were divided into six groups (female and male mice in each): six mice in each group were individually housed in cages. They were treated with distilled water and 20 μg/ml acetamidofluorene, 50 μg/mL fenaminosulf, 20 μg/mL sodium azide, and dosed with 50 μg/mL and 100 μg/mL by oral administration, respectively.

Animals were fasted overnight before the experiment, and were administered with a single dose of the extraction of KLBMPTC10 (20 mg/kg)/vehicle and any change in weight measured every seven days with a count of the number of poisoned and dead mice on each day. These tests were performed in duplicate (37). At the end of the study, all the surviving animals were sacrificed for blood analysis. Blood samples were collected from the heart and blood analysis was conducted at the weekend of the first and second week (7 and 14 days) after treatment. The non-heparinised blood was allowed to coagulate and then centrifuged to obtain serum. The serum was assayed for alanine aminotransferase (ALT), aspartate aminotransferase (AST), urea, uric acid, and creatinine (35).

### Statistical analysis

The experiments were conducted in triplicate. Data were reported as the mean ± SD for triplicate determinations. ANOVA and Tukey’s tests were undertaken to identify differences among means. Statistical significance was set at *P* < 0.05, and the results were expressed as mean ± SD.

## Results

### Isolation and identification of endophytic bacteria

In this study, a total of 31 isolate strains were identified from the inner tissues of *Taxus chinensis var. mairei*. The 31 isolates included 18 (KLBMPTC01-KLBMPTC18), eight (KLBMPTC19-KLBMPTC26), and five (KLBMPTC27-KLBMPTC31) strains from root, stem, and leaf tissue segments, respectively. In root specimens, 18 isolates were identified as *Brevibacillus* (one strain), *Viridibacillus* (one strain), and *Bacillus* (16 strains). In stem specimens, eight strains were distributed among *Microbacterium* (one strain), *Xanthomonas* (one strain), *Paracoccus* (one strain), and *Bacillus* (five strains) respectively. In leaf specimens, the five isolates belonged to *Bacillus* (four strains) and *Sphingomonas* (one strain). The specific information of the isolates is shown in Table 1 and their taxonomic status is shown in Figure 1.

**Figure 1.**
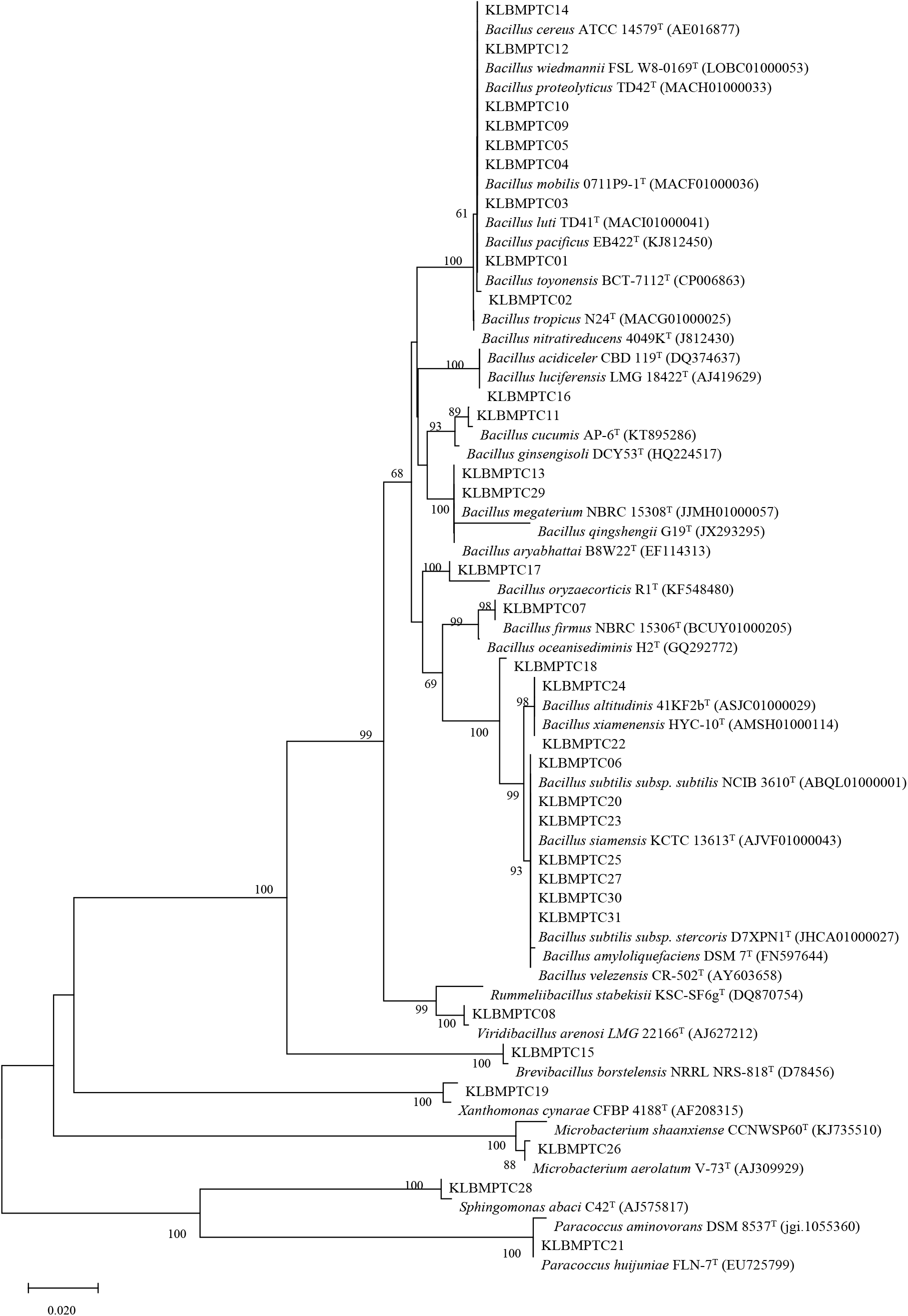
Phylogenetic neighbour-joining tree of the bacteria based on 16S rRNA gene sequences (numbers on branch nodes are bootstrap values: 1000 resampling operations; bar: 0.2% sequence divergence).

**Table 1.**
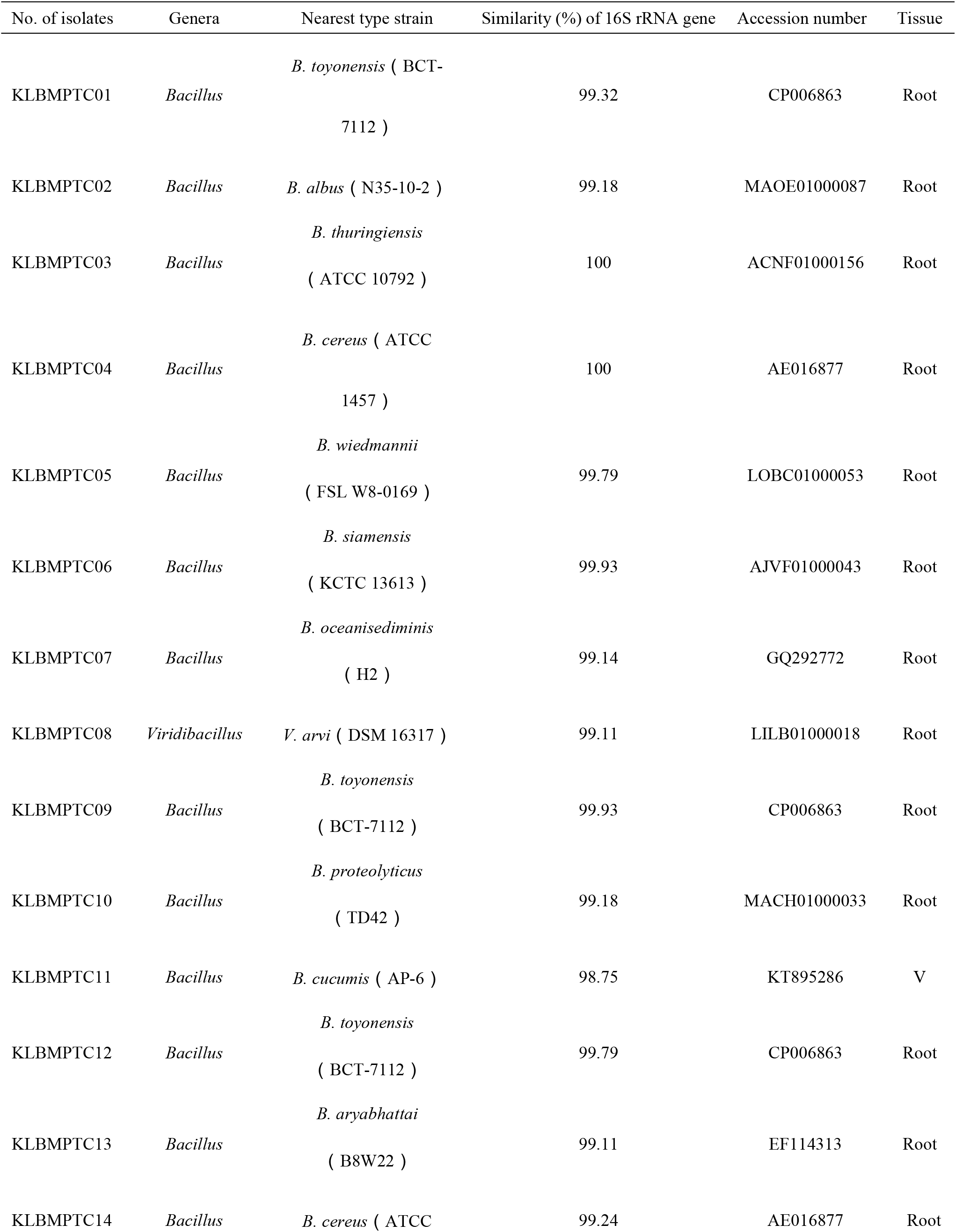
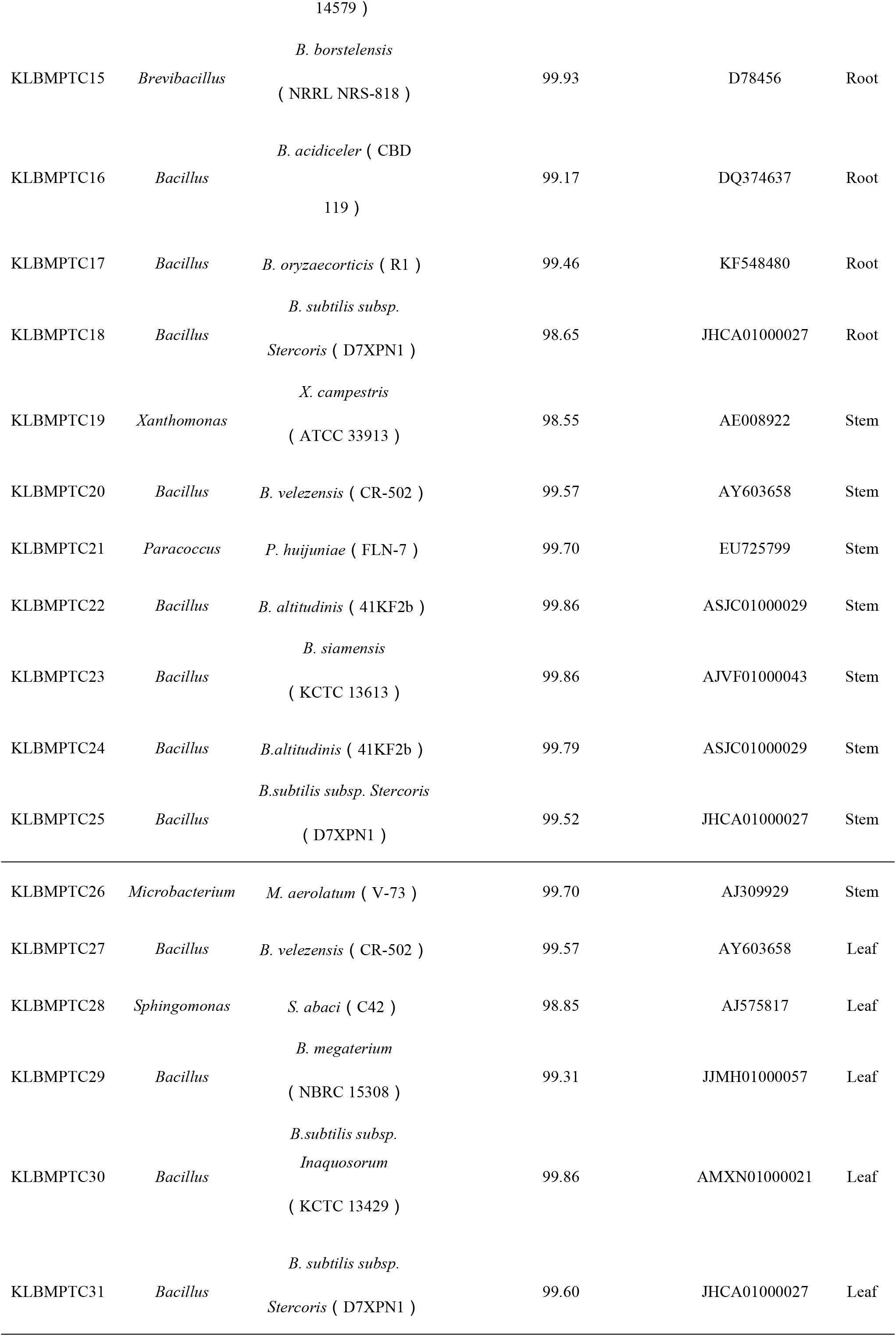
Identification endophytic bacteria by 16S rRNA gene sequences

### IAA-production and lipopeptide-production properties of the isolates

Biosynthesis of IAA showed differences among strains (Figure 2): 20 strains could produce IAA at between 0.2869 ± 0.02 ppm to 210.3955 ± 0.81 ppm. Lipopeptide-producing substances of these 20 strains were studied by LC-TOF-MS. Assay of the target lipopeptides returned the screening results listed in Table 2 and the relative content of the molecular ion peak intensity was as shown in Figure 3. According to Figure 3, the species and content had significant differences in lipopeptide compounds in IAA-producing bacteria. And the strain KLBMPTC10 contained the most abundant lipopeptide compounds among all tested IAA-producing bacteria, and its lipopeptide content was higher than that in other strains, followed by strains KLBMPTC19, KLBMPTC01, KLBMPTC02, and KLBMPTC29 which these strains all belonging to the genus *Bacillus*.

**Figure 2.**
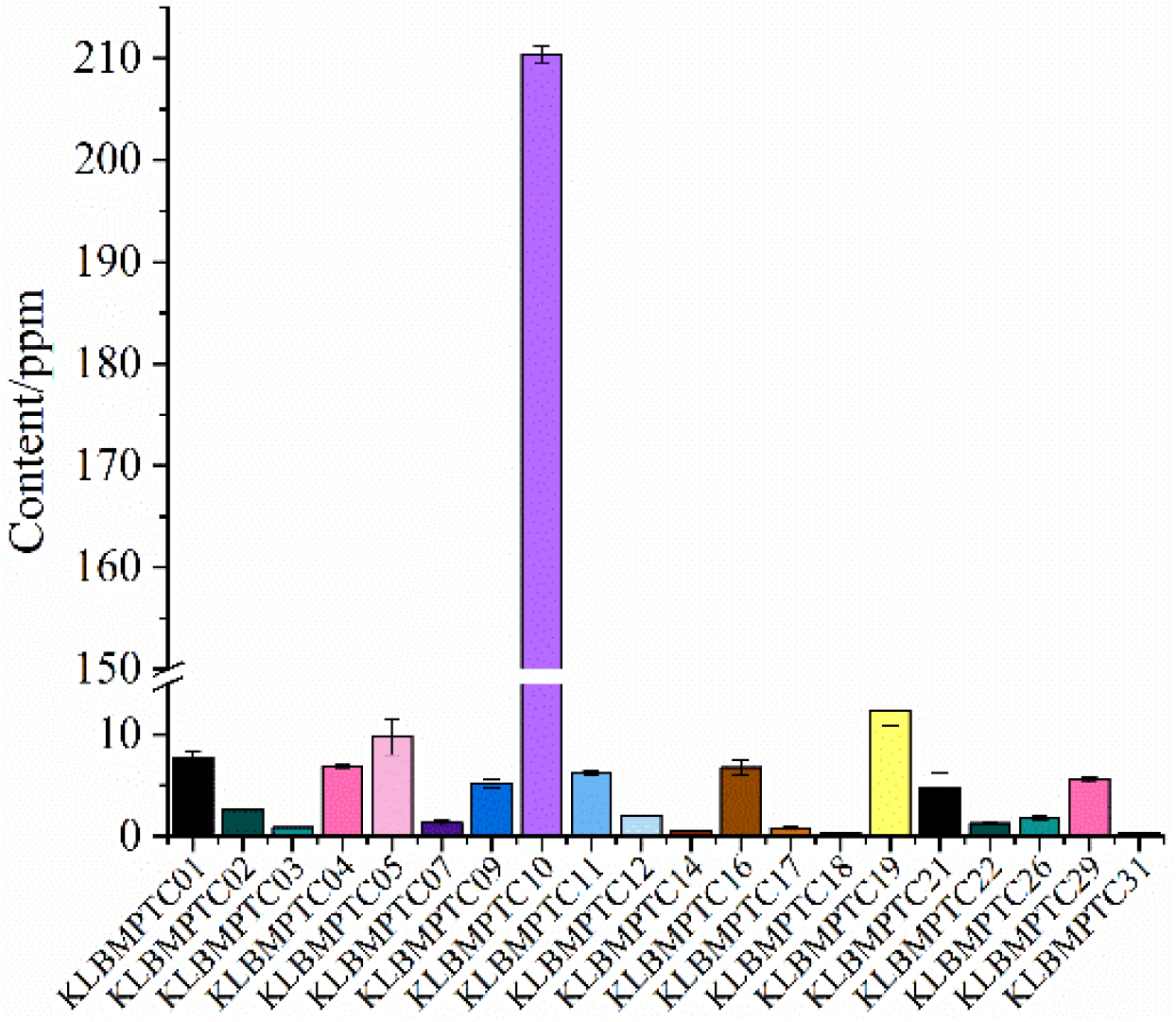
IAA-producing strains and contents thereof

**Figure 3.**
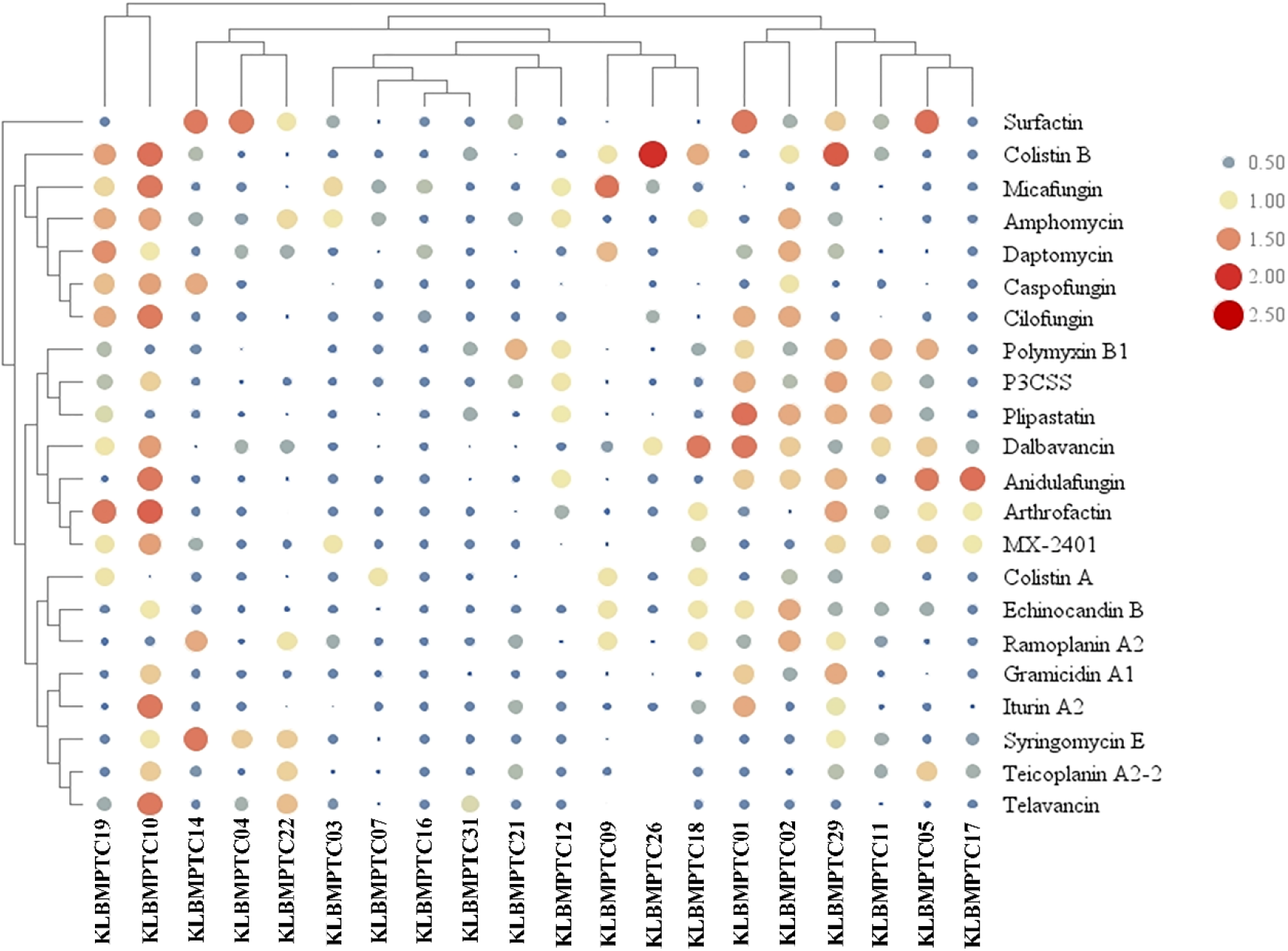
Cluster analysis of lipopeptide compounds produced by endophytic bacteria of *Taxus chinensis*

**Table 2.** Mass information: targeted lipopeptides

| No. | Compound | Predicted chemical formulae | Ion forms | Detected molecular weight (Da) | Theoretical molecular weight (Da) |
| --- | --- | --- | --- | --- | --- |
| 1 | Amphomycin | C<sub>58</sub>H<sub>91</sub>N<sub>13</sub>O<sub>20</sub> | [M+H]<sup>+</sup> | 1290.4184 | 1289.6497 |
| 2 | Anidulafungin | C<sub>58</sub>H<sub>73</sub>N<sub>7</sub>O<sub>17</sub> | [M+H]<sup>+</sup> | 1141.2371 | 1139.5057 |
| 3 | Arthrofactin | C<sub>64</sub>H<sub>111</sub>N<sub>11</sub>O<sub>20</sub> | [M+H]<sup>+</sup> | 1354.6328 | 1353.8001 |
| 4 | Caspofungin | C<sub>52</sub>H<sub>88</sub>N<sub>10</sub>O<sub>15</sub> | [M+H]<sup>+</sup> | 1093.3114 | 1092.6425 |
| 5 | Cilofungin | C<sub>49</sub>H<sub>71</sub>N<sub>7</sub>O<sub>17</sub> | [M+H]<sup>+</sup> | 1030.1203 | 1029.4901 |
| 6 | Colistin A | C<sub>53</sub>H<sub>100</sub>N<sub>16</sub>O<sub>13</sub> | [M+H]<sup>+</sup> | 1169.4698 | 1168.7650 |
| 7 | Colistin B | C<sub>52</sub>H<sub>98</sub>N<sub>16</sub>O<sub>13</sub> | [M+H]<sup>+</sup> | 1155.4325 | 1154.7493 |
| 8 | Dalbavancin | C<sub>88</sub>H<sub>100</sub>Cl<sub>2</sub>N<sub>10</sub>O<sub>28</sub> | [M+H]<sup>+</sup> | 1745.6929 | 1814.6080 |
| 9 | Daptomycin | C<sub>72</sub>H<sub>101</sub>N<sub>17</sub>O<sub>26</sub> | [M+H]<sup>+</sup> | 1620.6785 | 1619.7098 |
| 10 | Echinocandin B | C<sub>52</sub>H<sub>81</sub>N<sub>7</sub>O<sub>16</sub> | [M+H]<sup>+</sup> | 1060.2462 | 1059.57343 |
| 11 | Gramicidin A1 | C<sub>99</sub>H<sub>140</sub>N<sub>20</sub>O<sub>17</sub> | [M+H]<sup>+</sup> | 1882.2971 | 1881.06999 |
| 12 | Iturin A2 | C<sub>48</sub>H<sub>74</sub>N<sub>12</sub>O<sub>14</sub> | [M+H]<sup>+</sup> | 1043.1711 | 1042.54420 |
| 13 | Micafungin | C<sub>56</sub>H<sub>71</sub>N<sub>9</sub>O<sub>23</sub>S | [M+H]<sup>+</sup> | 1270.2786 | 1269.43780 |
| 14 | MX-2401 | C<sub>67</sub>H<sub>101</sub>N<sub>15</sub>O<sub>22</sub> | [M+H]<sup>+</sup> | 1468.6100 | 1467.72401 |
| 15 | P3CSS | C<sub>60</sub>H<sub>113</sub>N<sub>3</sub>O<sub>11</sub>S | [M+H]<sup>+</sup> | 1084.6226 | 1083.80904 |
| 16 | Plipastatin | C<sub>72</sub>H<sub>110</sub>N<sub>12</sub>O<sub>20</sub> | [M+H]<sup>+</sup> | 1463.7114 | 1462.79539 |
| 17 | Polymyxin B1 | C<sub>56</sub>H<sub>98</sub>N<sub>16</sub>O<sub>13</sub> | [M+H]<sup>+</sup> | 1203.4832 | 1202.74938 |
| 18 | Ramoplanin A2 | C<sub>119</sub>H<sub>154</sub>N<sub>21</sub>O<sub>40</sub> | [M+H]<sup>+</sup> | 2554.0796 | 2517.06565 |
| 19 | Surfactin | C<sub>53</sub>H<sub>93</sub>N<sub>7</sub>O<sub>13</sub> | [M+H]<sup>+</sup> | 1036.3408 | 1035.68259 |
| 20 | Syringomycin E | C<sub>53</sub>H<sub>85</sub>ClN<sub>14</sub>O<sub>17</sub> | [M+H]<sup>+</sup> | 1189.7872 | 1224.59002 |
| 21 | Teicoplanin A2-2 | C<sub>88</sub>H<sub>97</sub>ClN<sub>12</sub>O<sub>33</sub> | [M+H]<sup>+</sup> | 1850.2066 | 1884.59640 |
| 22 | Telavancin | C<sub>80</sub>H<sub>106</sub>Cl<sub>2</sub>N<sub>11</sub>O<sub>27</sub>P | [M+H]<sup>+</sup> | 1652.6407 | 1753.63688 |

At the same time, surfactin and iturin A2 in strains KLBMPTC10, KLBMPTC01, and KLBMPTC29 were quantified by LC-MS/MS. A patent strain *Bacillus amyloliquefaciens* CGMCC5569 obtained in previous laboratory work was used as a positive control strain: the results are shown in Figure 4. Compared to the control strain, it can be seen that three endophytic bacteria showed significant differences in the production of two typical lipopeptide compounds (surfactin and iturin A2) which were typical lipopeptide compounds in *Bacillus*. Among them, the surfactin-production ability of strain KLBMPTC29 was better than that of other two strains, the content of which was 24.47±4.05%. The ability of strain KLBMPTC10 to produce iturin A2 was greater than that of the other two strains, and the content thereof was 26.06± 2.68%.

**Figure 4.**
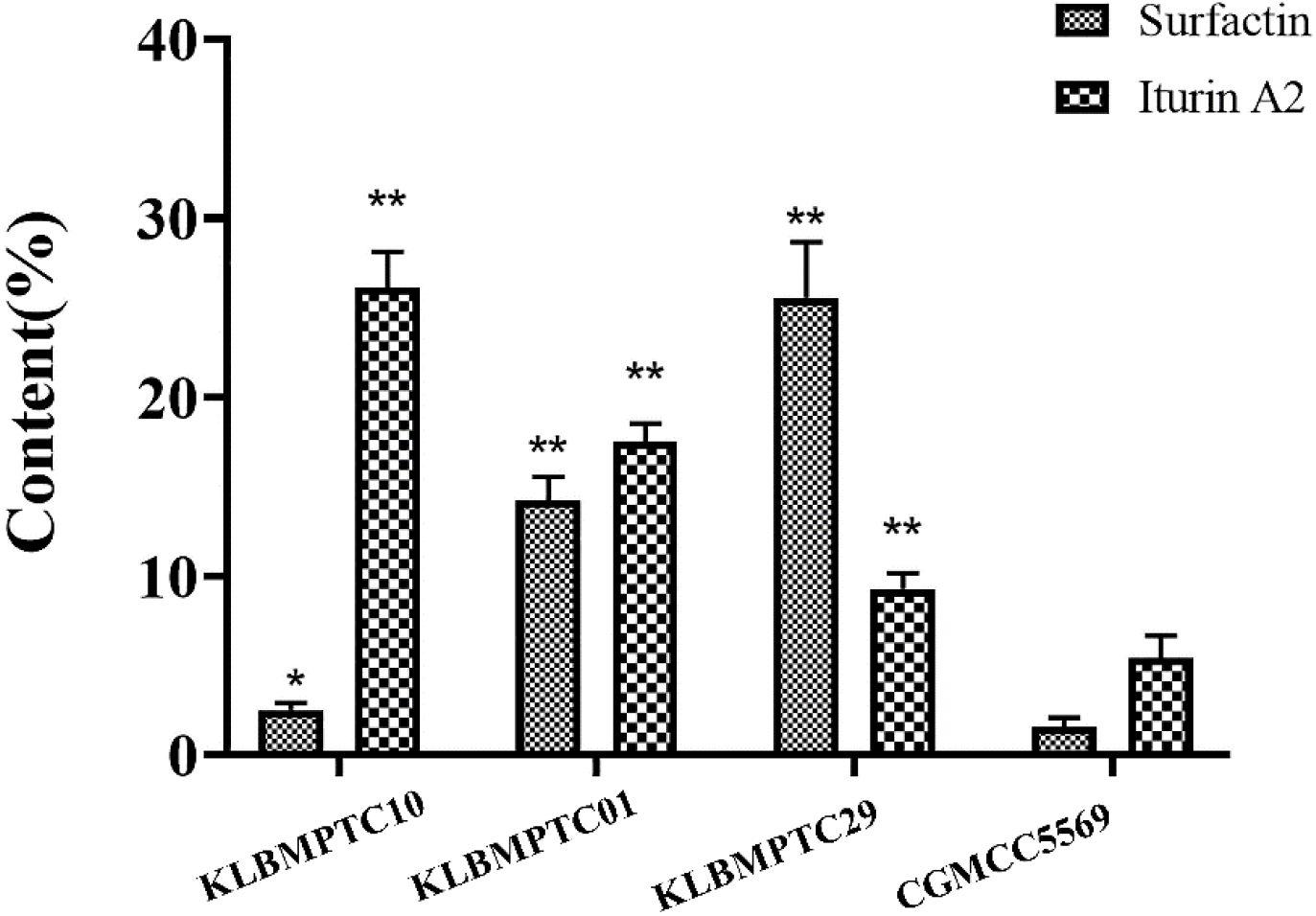
Quantitative calculation of two typical lipopeptides in three endophytic bacteria strains based on LC-MS/MS

### *In vivo* toxicity of KLBMPTC10 extraction on mice

According to the results presented in “Isolation and identification of endophytic bacteria” and “IAA-production and lipopeptide-production properties of the isolates”, we selected strain KLBMPTC10 to evaluate the safety of mice. Firstly, we observed the possible mutagenesis of KLBMPTC10 extraction by Ames test. The Ames test performed *in vitro*, in the short-term to evaluate possible mutagenic effects caused by chemicals. The TA97, TA98, TA100, and TA102 strains of *S. typhimurium* used in the Ames test carry a defective gene (a mutant) that prevent them from synthesising histidine: however, when there are mutagenic chemical compounds present, defective genes can be mutated back to produce histidine by reversing the mutation, so they can grow on a histidine-free plate. Usually, the number of colonies reverting spontaneously is quasi-stable: if the bacterial extract of strain KLBMPTC10 induces back-mutation colonies that exceed twice the number of spontaneous back-mutations, it is considered to be mutagenic (41). As the result was shown in Table 3, under the treatment of bacterial extracts of different concentrations of strain KLBMPTC10, the number of colonies induced by TA97, TA98, TA100, and TA102 strains of *S. typhimurium* did not exceed twice the number of spontaneous reverse mutations. Therefore, the bacterial extract of bacteria KLBMPTC10 was not mutagenic.

**Table 3.** Ames testing results

| Groups | TA97 | TA98 | TA100 | TA102 |
| --- | --- | --- | --- | --- |
| Blank control | 114 ± 15 | 117 ± 12 | 149 ± 17 | 214 ± 15 |
| Solvent | 109 ± 20 | 114 ± 13 | 152 ± 8 | 224 ± 16 |
| Positive control |  |  |  |  |
| 2-Acetamidofluorene<br>(20 µg/mL) | 1455 ± 86 | 1277 ± 62 | 542 ± 14 | 814 ± 14 |
| Dexon<br>(50 µg/mL) Dexon<br>(50 µg/mL) | 1865 ± 160 | 1508 ± 94 | 614 ± 9 | 692 ± 10 |
| Sodium azide<br>(20 µg/mL) Sodium<br>azide<br>(20 µg/mL) | 997 ± 142 | 1022 ± 97 | 529 ± 11 | 542 ± 9 |
| The bacterial<br>extract of strain<br>KLBMPTC10 |  |  |  |  |
| 5 µg/mL | 117 ± 25 | 114 ± 25 | 134 ± 9 | 211 ± 18 |
| 10 µg/mL | 122 ± 13 | 125 ± 17 | 142 ± 10 | 254 ± 16 |
| 20 µg/mL | 147 ± 10 | 127 ± 35 | 174 ± 14 | 248 ± 11 |
| 100 µg/mL | 151 ± 12 | 124 ± 14 | 150 ± 5 | 262 ± 10 |

After the two-week extraction treatment, there was no mortality, unusual changes in behaviour, locomotive activity, or signs of intoxication observed. The body mass of mice treated for 0 days, 7 days, and 14 days is shown in Table 4. No significant differences were found in the body mass of six groups. Simultaneously, alanine aminotransferase (ALT), aspartate aminotransferase (AST), urea, uric acid, and creatinine were measured in blood sampled from the heart of the mice under test. The data from the blood analyses are shown in Figures 5 and 6. According to the results, compared with positive control, there were no significant differences in ALT, AST, urea, uric acid, and creatinine on blank control and the other two treatment of bacterial extracts of strain KLBMPTC10 groups. And these all indicated that the bacterial extracts of strain KLBMPTC10 had no harmful effect on mice.

**Figure 5.**
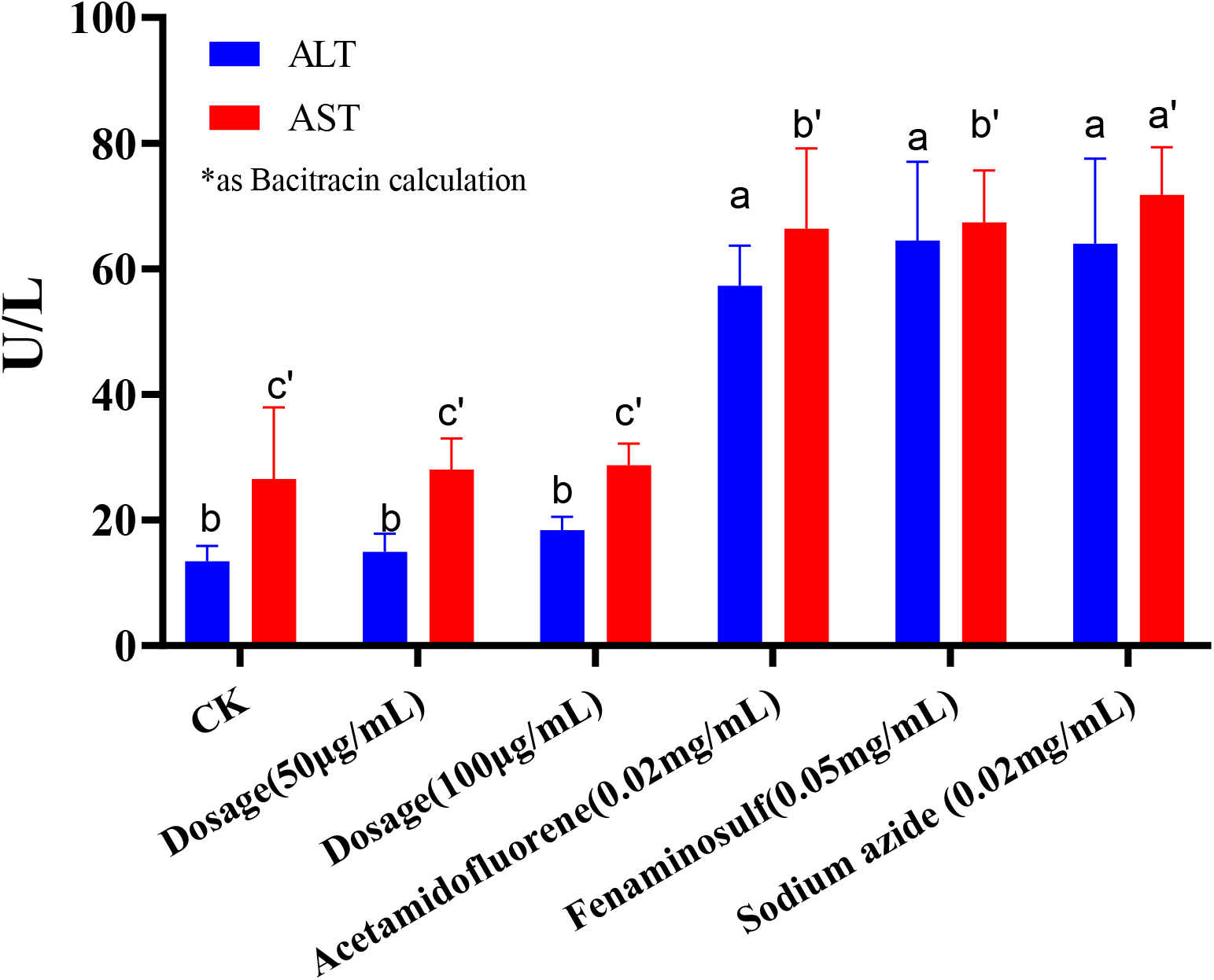
the changes of the ALT and AST under the treatment

**Figure 6.**
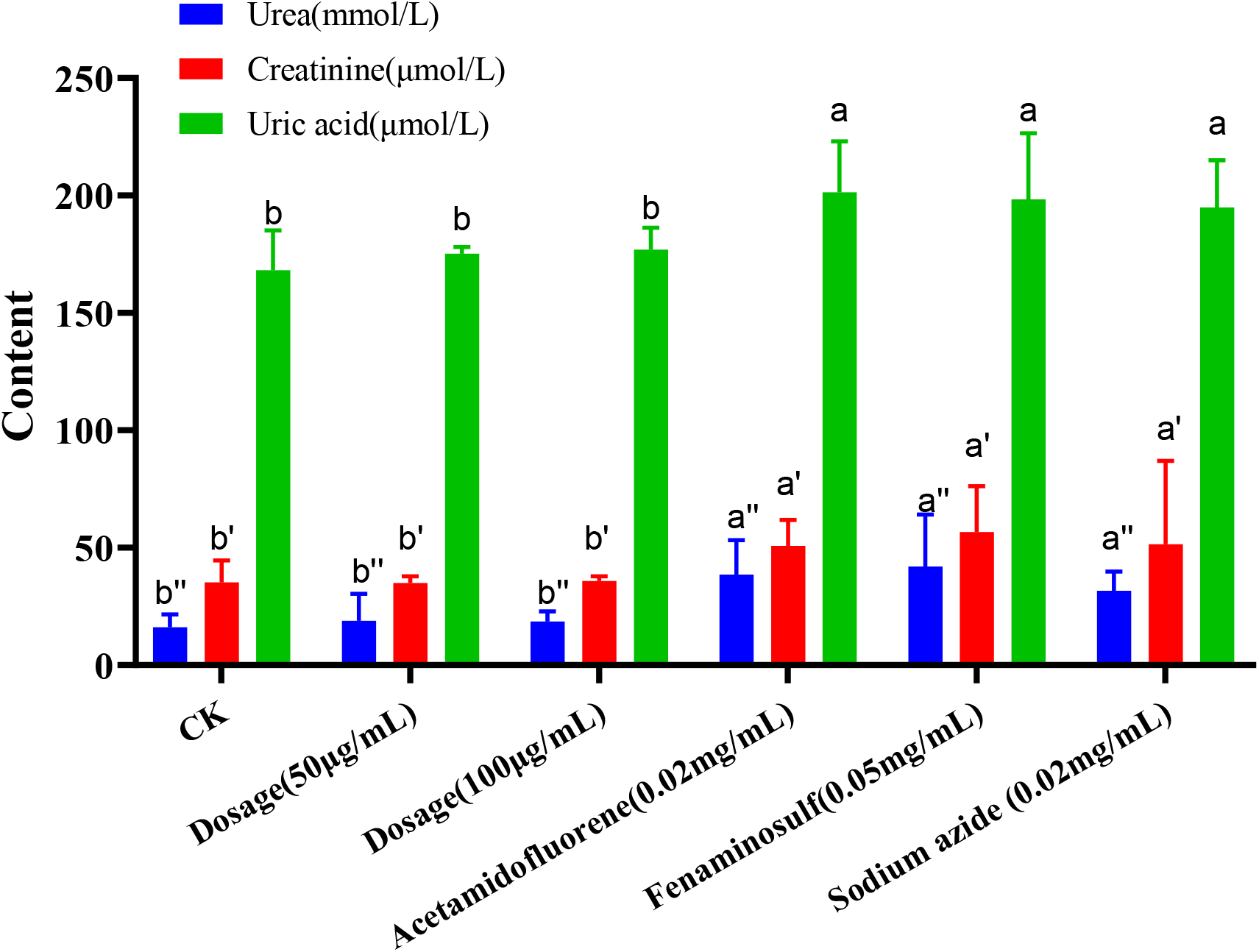
the results of the Urea and Creatinine and Uric acid in Blood analysis

**Table 4.** Body mass of mice treated for 0 days, 7 days, and 14 days

| Group | 0 days | 7 days | 14 days |
| --- | --- | --- | --- |
| Male |  |  |  |
| CK | 16.25 ± 0.54 | 22.63 ± 1.25 | 29.70 ± 1.22 |
| 50 µg/mL | 15.97 ± 0.42 | 21.44 ± 0.47 | 27.33 ± 0.84 |
| 100 µg/mL | 16.29 ± 0.44 | 20.66 ± 0.42 | 25.08 ± 1.14 |
| Female |  |  |  |
| CK | 16.47 ± 0.77 | 24.04 ± 0.92 | 30.01 ± 0.97 |
| 50 µg/mL | 16.02 ± 0.73 | 23.11 ± 0.88 | 28.29 ± 0.75 |
| 100 µg/mL | 17.07 ± 0.54 | 22.93 ± 0.58 | 26.38 ± 0.73 |

## Discussion

It is understood that 1/3 of the world’s soil is in a severely damaged state (42) and this affects the growth of plants. And the reason is that the environmental pollution caused by the development of modern science and technology occupies a large part (43). In agriculture, the environmental pollution mainly refers to excessive use of chemical fertilizers, pesticides and heavy metal which not only has a negative effect on the environment, but also on the reproductive and physiological functions of humans and animals (44, 45). Therefore, people hope to mitigate the pollution to the soil by looking for environmentally friendly methods, and the microorganisms may be a better choice (16). Studies have shown that the overwhelming majority of *Bacillus* can not only promote plant growth, but also produce certain antimicrobial and insecticidal substances for biological control (10, 16).

IAA is a growth-promoting substance which can promote the plant growth and stress resistance(46). It plays an important role in plant life cycle. Surfactin, iturin, and fengycin are the main lipopeptide active substances produced by *Bacillus* sp. These substances have shown inhibitory effects on a variety of pathogens and pest whether in plate experiments or plant experiments (47, 48). In this study, we first isolated endophytic bacteria from *Taxus chinensis* and identified them with 16S rDNA, and then we screened the IAA producing ability of these strains. The results showed that we isolated a high IAA producing *Bacillus* from the roots of *Taxus chinesis*. Its IAA production in testing conditions is higher than other *Bacillus* strains(49, 50). In addition, we used HPLC-TOFMS to test the ability the ability of these IAA producing strains to produce lipopeptide compounds which may help reduce the damage of some diseases or insect pests and is environmentally friendly. And we found that KLBMPTC10 is a high lipopeptide producer strain and the quantitative results showed that the ability of producing iturin A2 was better than other strains, and the content was 26.06 ± 2.68%. And iturin compounds have significant inhibitory activity on fungal growth and it has been widely used in agriculture (51).

In addition, microbes are harmless to humans and animals, which is the most important evaluation index to evaluate their potential application. So according to the above characteristics evaluation results of strain KLBMPTC10, and to further understand its potential value, we also undertook Ames and acute oral toxicity, testing in mice to evaluate its safety when used on mammals. Ames test can evaluate possible mutagenic effects caused by chemicals in vitro in short term (52). We can consider it as a mutagen if bacterial extract of strain KLBMPTC10 induces back-mutation colonies that exceed twice the number of spontaneous back-mutations (41). ALT, AST, urea, uric acid data in mice can assess whether liver and kidney functions have been damaged (53). In this study, the results of Ames test and acute oral toxicity bacterial extract of strain KLBMPTC10 has not mutagenicity and non-toxic to mice. And this shows that the strain KLBMPTC10 has great potential value in application to forest resource protection and biofertiliser production.

## Funding information

This research was supported by the National Natural Science Foundation of China (Grant no. 31770613, 31571735, 81522049, 31800327), the Research Innovation Program for College Graduates of Jiangsu Province (Grant no.771241822), Zhejiang Provincial Ten-thousand Program for Leading Talents of Science and Technology Innovation (2018R52050), Zhejiang Provincial Program for the Cultivation of High-level Innovative Health talents, and Opening project of Zhejiang provincial preponderant and characteristic subject of Key Laboratory, Zhejiang Chinese Medical University (ZYAOX2018004).

## Conflict of interest

The authors declare no competing financial interest.

